# Scanning the Horizon: Towards transparent and reproducible neuroimaging research

**DOI:** 10.1101/059188

**Authors:** Russell A. Poldrack, Chris I. Baker, Joke Durnez, Krzysztof J. Gorgolewski, Paul M. Matthews, Marcus Munafò, Thomas E. Nichols, Jean-Baptiste Poline, Edward Vul, Tal Yarkoni

## Abstract

Functional neuroimaging techniques have transformed our ability to probe the neurobiological basis of behaviour and are increasingly being applied by the wider neuroscience community. However, concerns have recently been raised that the conclusions drawn from some human neuroimaging studies are either spurious or not generalizable. Problems such as low statistical power, flexibility in data analysis, software errors, and lack of direct replication apply to many fields, but perhaps particularly to fMRI. Here we discuss these problems, outline current and suggested best practices, and describe how we think the field should evolve to produce the most meaningful answers to neuroscientific questions.

## Main text

Neuroimaging, particularly using functional magnetic resonance imaging (fMRI), has become the primary tool of human neuroscience^1^, and recent advances in the acquisition and analysis of fMRI data have provided increasingly powerful means to dissect brain function. The most common form of fMRI (known as “blood oxygen level dependent” or BOLD fMRI) measures brain activity indirectly through localized changes in blood oxygenation that occur in relation to synaptic signaling^2^. These signal changes provide the ability to map activation in relation to specific mental processes, identify functionally connected networks from resting fMRI^3^, characterize neural representational spaces^4^, and decode or predict mental function from brain activity^5,6^. These advances promise to offer important insights into the workings of the human brain, but also generate the potential for a “perfect storm” of irreproducible results. In particular, the high dimensionality of fMRI data, relatively low power of most fMRI studies, and the great amount of flexibility in data analysis all potentially contribute to a high degree of false positive findings.

Recent years have seen intense interest in the reproducibility of scientific results and the degree to which some problematic yet common research practices may be responsible for high rates of false findings in the scientific literature, particularly within psychology but also more generally^7–9^. There is growing interest in “meta-research”^10^, and a corresponding growth in studies investigating factors that contribute to poor reproducibility. These factors include study design characteristics which may introduce bias, low statistical power, and flexibility in data collection, analysis, and reporting — termed “researcher degrees of freedom” by Simmons and colleagues^8^. There is clearly concern that these issues may be undermining the value of science – in the UK, the Academy of Medical Sciences recently convened a joint meeting with a number of other funders to explore these issues, while in the US the National Institutes of Health has an ongoing initiative to improve research reproducibility^11^.

In this article we outline a number of potentially problematic research practices in neuroimaging that can lead to increased risk of false or exaggerated results. For each problematic research practice, we propose a set of solutions. While most proposed solutions are uncontroversial in principle, their implementation is often challenging for the research community and best practices are not necessarily followed. Many of these solutions arise from the experience of other fields with similar problems (particularly those dealing with similarly large and complex data sets, such as genetics; Box 1). We would note that while our discussion here focuses on functional MRI, many of the same issues are relevant for other types of neuroimaging, such as structural or diffusion MRI.

### Statistical power

The analyses of Button and colleagues^12^ provided a wake-up call regarding statistical power in neuroscience, particularly by highlighting the point (raised earlier by Ioannidis^7^) that low power not only reduces the likelihood of finding a true result if it exists, but also raises the likelihood that any positive result is false, as well as causing substantial inflation of observed positive effect sizes^13^. In the context of neuroimaging, Button and colleagues considered only structural MRI studies. In order to assess the current state of statistical power in fMRI studies, we performed an analysis of sample sizes and the resulting statistical power of fMRI studies over the past 20 years.

To gain a perspective on how sample sizes have changed over this time period, we obtained sample sizes from fMRI studies using two sources. First, manually annotated sample size data for 583 studies were obtained from published meta-analyses^14^. Second, sample sizes were automatically extracted from the Neurosynth database^15^ for 548 studies published between 2011 and 2015 (by searching for regular expressions reflecting sample size, e.g. “13 subjects”, “n=24”) and then manually annotated to confirm automatic estimates and identify single-group versus multiple-group studies. (Data and code to generate all figures in this paper are available from the Open Science Framework at https://osf.io/spr9a/.) Figure 1a shows that sample sizes have steadily increased over the past two decades, with the median estimated sample size for a single-group fMRI study in 2015 at 28.5. A particularly encouraging finding from this analysis is that the number of recent studies with large samples (greater than 100) is rapidly increasing (from 8 in 2012 to 17 in 2015, in the studied sample), suggesting that the field may be progressing towards adequately powered research. On the other hand, the median group size in 2015 for fMRI studies with multiple groups was 19 subjects, which is below even the absolute minimum sample size of 20 per cell proposed by Simonsohn et al.^8^.

**Figure 1.**
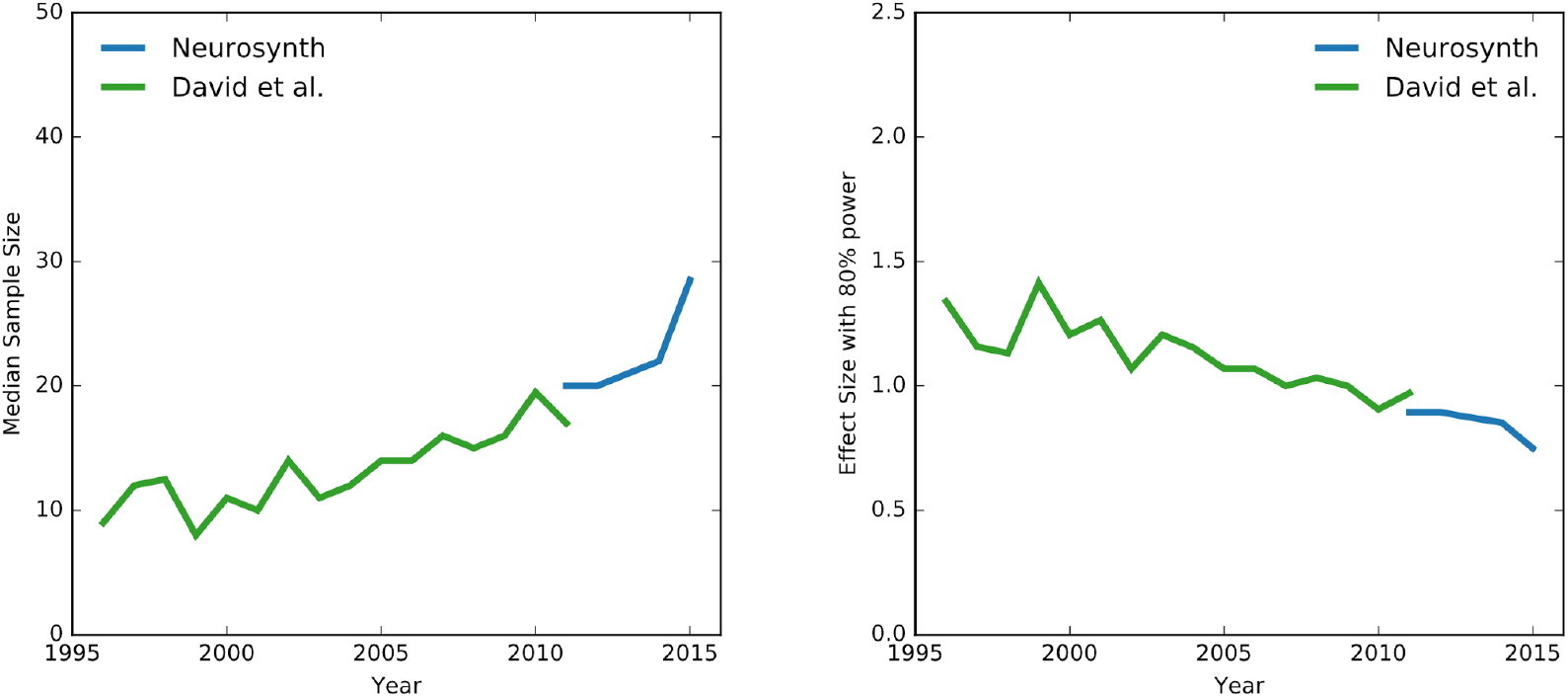
Sample size estimates and estimated power for fMRI studies. a | 1131 sample sizes over more than 20 years obtained from two sources: 583 sample sizes by manual extraction from published meta-analyses by David et al.^14^, and 548 sample sizes obtained by automated extraction from the Neurosynth database^15^ with manual verification. These data demonstrate that sample sizes have steadily increased over the last two decades, with a median estimated sample size of 28.5 as of 2015. b | Using the sample sizes from the left panel, we estimated the standardized effect size required to detect an effect with 80% power for a whole-brain linear mixed-effects analysis using a voxelwise 5% familywise error rate threshold from random field theory^16^ (see main text for details). Median effect size for which studies were powered to find in 2015 was 0.75. Data and code to generate these figures are available at https://osf.io/spr9a/; see Supplementary materials for a version with all datapoints depicted.

In order to assess the implications of these results for statistical power, for each of the 1131 sample sizes shown in Figure 1a we estimated the standardized effect size that would be required to detect an effect with 80% power (the standard level of power for most fields) for a whole-brain linear mixed-effects analysis using a voxelwise 5% familywise error (FWE) rate threshold from random field theory^16^ (a standard thresholding level for neuroimaging studies). In other words, we found the minimum effect size that would have been needed in each of these studies in order for the difference to be considered statistically significant with an 80% probability, given the sample size. We quantify the standardised effect size using Cohen’s D, computed as the average effect divided by the standard deviation for the data.

To do this, we assumed that each study used a statistical map with T-values in an MNI152 (Montreal Neurological Institute) template space with smoothness of three times the voxel size (full width at half maximum), a commonly used value for smoothness in fMRI analysis. The MNI152 template is a freely available template, obtained from an average T1 scan for 152 subjects with a resolution of 2 millimeters and a volume within the brain mask of 228483 voxels, used by default in most fMRI analysis software. We assume that in each case there would be one active region, with voxelwise standardised effect size D; that is, we assume that for each subject, all voxels in the active region are on average D standardised units higher in their activity than the voxels in the non-active region, and that the active region is 1,600 mm^2^ (200 voxels). To calculate the voxelwise statistical significance threshold for the active region in this model statistical map, we used the function *ptoz* from the FSL^17^ software package, which computes a FWE threshold for a given volume and smoothness using the Euler Characteristic derived from Gaussian random field theory ^18^. This approach ensures that the probability of a voxel in the non-active brain region exceeding this significance threshold is controlled at 5%; the resulting significance threshold, t_α_, is equal to 5.12.

The statistical power is defined as the probability that the local maximum peak of activation in the active region exceeds this significance threshold. This probability was computed using a shifted version of the null local maximum distribution, with shift of D*sqrt(n) to reflect a given effect size and sample size. The median effect size needed to exceed the significance threshold in each of the studies was found by selecting the effect size D that results in statistical power higher than 0.80 as computed in the previous step.

Figure 1b shows the median effect sizes needed to establish significance, with 80% power and alpha = 0.05. Despite the decreases in these hypothetical required effect sizes over the past 20 years, Fig. 1b shows that in 2015 the median study is only sufficiently powered to detect relatively large effects of greater than ~0.75. Given that many of the studies will be assessing group differences or brain activity–behaviour correlations (which will inherently have lower power than average group activation effects), this represents an optimistic lower bound on the powered effect size.

Indeed, the analysis presented in Box 2 demonstrates that typical effect sizes observed in task-related BOLD imaging studies fall well below this level. Briefly, we analysed BOLD data from 186 individuals who were imaged using fMRI while performing motor, emotion, working memory and gambling tasks as part of the Human Connectome Project^19^. Assessing effect sizes in fMRI requires the definition of an independent region of interest that captures the expected area of activation within which the effect size can be measured. While there are a number of approaches to defining regions ^20,21^, we created masks defined by the intersection between functional activation (identified from Neurosynth.org as regions consistently active in studies examining the effects of ‘motor’, ‘emotion’, ‘gambling’ and ‘working memory’ tasks) and anatomical masks (defined using the Harvard–Oxford probabilistic atlas^22^, based on the published regions of interest from the HCP)^23^. Within these intersection masks, we then determined the average task-related increases in BOLD signal — and the effect size (Cohen’s D) — associated with each different task. Additional details are provided in Box 2. The figure in Box 2, which lists the resulting BOLD signal changes and inferred effect sizes, demonstrates that realistic effect sizes – i.e. BOLD changes associated with a range of cognitive tasks - in fMRI are surprisingly small: even for powerful tasks such as the motor task which evokes median signal changes of greater than 4%, 75% of the voxels in the masks have a standardised effect size smaller than 1. For more subtle tasks, such as gambling, only 10% of the voxels in our masks demonstrated standardised effect sizes larger than 0.5. Thus the average fMRI study remains poorly powered for capturing realistic effects, particularly given that the HCP data are of particularly high quality, and thus the present estimates are likely greater than what would be found with most standard fMRI datasets.

### Solutions

When possible, all sample sizes should be justified by an *a priori* power analysis. A number of tools are available to enable power analyses for fMRI (for example, neuropowertools.org (see Further information; described in ref ^24^) and fmripower.org (see Further information; described in ref. ^25^). However, one must be cautious in extrapolating effect sizes from small studies, because they are almost certain to be inflated. When previous data are not available to support a power analysis, one can instead identify the sample size that would support finding the minimum effect size that would be theoretically informative (e.g. based on results from Box 2). The use of heuristic sample size guidelines (for example, based on sample sizes used in previously published studies) is likely to result in a misuse of resources, either by collecting too many or (more likely) too few subjects.

The larger sample sizes that will result from use of power analysis will have significant implications for researchers: Given that research funding will likely not increase to accommodate these larger samples, this implies that fewer studies will be funded, and that researchers with fewer resources will have a more difficult time performing research that meets these standards. This will hit trainees and junior researchers particularly hard, and the community needs to develop ways to address this challenge. We do not believe that the solution is to admit weakly powered studies simply because of a lack of resources. This situation is in many ways similar to the one faced in the field of genetics, which realized more than a decade ago that weakly-powered genetic association studies were unreliable, and moved to the use of much larger samples with high power to detect even very small associations and as well as the enforcement of replication. This has been accomplished through the development of large-scale consortia, which have amassed samples in the tens or hundreds of thousands (see Box 1). There are examples of successful consortia in neuroimaging, including the 1000 Functional Connectomes Project and its International Neuroimaging Data-sharing Initiative (INDI)^3,26^ and the ENIGMA (Enhancing Neuro Imaging Genetics by Meta-Analysis) consortium^27^. With such consortia come inevitable challenges of authorship and credit^28^, but here again we can look to other areas of research that have met these challenges.

In some cases, researchers must necessarily use an insufficient sample size in a study, due to limitations in the specific sample (for example, when studying a rare patient group). In such cases, there are three commonly used options to improve power. First, researchers can choose to collect a much larger amount of data from each individual, and present results at the individual level rather than at the group level^29,30^--though the resulting inferences cannot then be generalized to the population as a whole. Second, researchers can use more liberal statistical thresholding procedures, such as methods controlling the false discovery rate (FDR). However, it should be noted that the resulting higher power comes at the expense of more false positive results and should therefore be used with caution; any results must be presented with the caveat that they have an inflated false positive rate. Third, researchers may restrict the search space using a small number of *a priori* regions of interest (ROIs) or an independent ‘functional localizer’ (a separate scan used to identify regions based on their functional response, such as retinotopic visual areas or face-responsive regions) to identify specific ROIs for each individual. It is essential that these ROIs (or a specific functional localizer strategy) be explicitly defined before any analyses. This is important because it is always possible to develop a *post hoc* justification for any specific ROI on the basis of previously published papers — a strategy that results in an ROI that appears independent but actually has a circular definition and thus leads to meaningless statistics and inflated Type I errors. By analogy to the idea of HARKing (hypothesizing after results are known; in which the results of exploratory analyses are presented as having been hypothesized from the beginning)^31^, we refer to this as SHARKing (selecting hypothesized areas after results are known). We would only recommend the use of restricted search spaces if the exact ROIs and hypotheses are pre-registered^32,33^.

Finally, we note the potential for Bayesian methods to make the best use of small, underpowered samples. These approaches stabilize low-information estimates, shrinking them towards anticipated values characterized by prior distributions. While Bayesian methods have not been widely used in the whole-brain setting due to the computational challenge of specifying a joint model over all voxels, GPU’s may provide the acceleration needed to make these methods practical (e.g.^34^). These methods also require the specification of priors, which often remains a challenge.

### Problem: Flexibility in data analysis

The typical fMRI analysis workflow contains a large number of preprocessing and analysis operations, each with choices to be made about parameters and/or methods (see Box 3). Carp^35^ applied 6,912 analysis workflows (using the SPM^36^ and AFNI^37^ software packages) to a single data set and quantified the variability in resulting statistical maps. This revealed that some brain regions exhibited more substantial variation across the different workflows than did other regions. This issue is not unique to fMRI; for example, similar issues have been raised in genetics^38^. These “researcher degrees of freedom” can lead to substantial inflation of Type I error rates^8^ whenever an analysis decision is made based partly on the observed data--even when there is no intentional “p-hacking”, and only a single analysis is ever conducted^9^.

Exploration is key to scientific discovery, but rarely does a research paper comprehensively describe the actual process of exploration that led to the ultimate result; to do so would render the resulting narrative far too complex and murky. As a clean and simple narrative has become an essential component of publication, the intellectual journey of the research is often obscured. Instead, reports often engage in HARKing^31^. Because HARKing hides the number of data-driven choices made during analysis, it can strongly overstate the actual evidence for a hypothesis. There is arguably a great need to support the publication of exploratory studies without forcing those studies to masquerade as hypothesis-driven science, while at the same time realizing that such exploratory findings (like all scientific results) will ultimately require further validation in independent studies.

### Solutions

We recommend pre-registration of methods and analysis plans. The details to be pre-registered should include planned sample size, specific analysis tools to be used, specification of predicted outcomes, and definition of any ROIs that will be used for analysis. The Open Science Framework (http://osf.io) and AsPredicted.org provide established platforms for pre-registration; the former allows an embargo period in which the registration remains private, obviating some concerns about ideas being disclosed while still under investigation. In addition, some journals now provide the ability to submit a “Registered Report”, in which the hypotheses and methods are reviewed prior to data collection, and the study is guaranteed publication regardless of the outcome^39^; see ^40,41^ for examples of such reports, and https://osf.io/8mpji/wiki/home/ for a list of journals offering the Registered Report format. Exploratory analyses (including any deviations from planned analyses) should be clearly distinguished from planned analyses in the publication. Ideally, results from exploratory analyses should be confirmed in an independent validation data set.

While there are concerns regarding the degree to which flexibility in data analysis may result in inflated error rates, we do *not* believe that the solution should be to constrain researchers by specifying a particular set of methods that must be used. Many of the most interesting findings in fMRI have come from the use of novel analysis methods, and we do not believe that there will be a single best workflow for all studies; in fact, there is direct evidence that different studies or individuals will likely benefit from different workflows ^42^. We believe that the best solution is to allow flexibility, but require that all exploratory analyses be clearly labeled as such, and strongly encourage validation of exploratory results (e.g. through the use of a separate validation dataset).

### Problem: Multiple comparisons

The most common approach to neuroimaging analysis involves “mass univariate” testing in which a separate hypothesis test is performed for each voxel. In such an approach, the false positive rate will be inflated if there is no correction for multiple tests. A humorous example of this was seen in the now-infamous “dead salmon” study reported by Bennett and colleagues^43^, in which “activation” was detected in the brain of a dead salmon, which disappeared when the proper corrections for multiple comparisons were performed.

Figure 2 presents a similar example in which random data can be analysed (incorrectly) to lead to seemingly impressive results, through a combination of failure to adequately correct for multiple comparisons and circular ROI analysis. We generated random simulated fMRI and behavioral data from a Gaussian distribution (mean±standard deviation = 1000±100 for fMRI data, 100±1 for behavioral data) for 28 simulated subjects (based on the median sample size found in the analysis of Figure 1 for studies from 2015). For the fMRI data, we simulated statistical values at each voxel for a comparison of activation and baseline conditions for each of the simulated subjects within the standard MNI152 mask, and then spatially smoothed the image with a 6mm Gaussian kernel, based on the common smoothing level of 3 times the voxel size. A univariate analysis was performed using FSL to assess the correlation between activation in each voxel and the simulated behavioural regressor across subjects, and the resulting statistical map was thresholded at p < 0.001 and with a 10-voxel extent threshold (which is a common heuristic correction shown by Eklund et al.^44^ to result in highly inflated levels of false positives). This approach revealed a cluster of false positive activation in the superior temporal cortex in which the simulated fMRI data are highly correlated with the simulated behavioural regressor (Fig. 2a).

**Figure 2:**
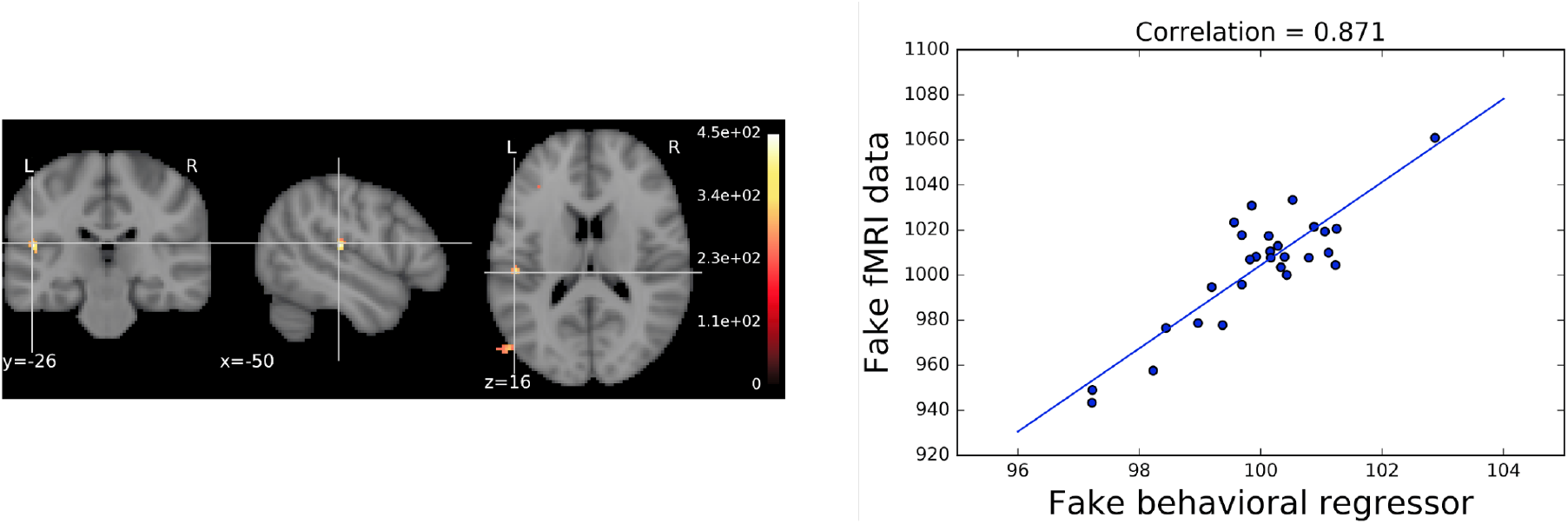
Small samples, uncorrected statistics and circularity can produce misleadingly large effects. Seemingly impressive brain-behavior association can arise from completely random data through the use of uncorrected statistics and circular ROI analysis to capitalize on the large sampling error arising from small samples. With the informal P<0.001 and cluster size k>10 thresholding, the analysis revealed a cluster in the superior temporal gyrus (left panel); signal extracted from that cluster (i.e., using circular analysis) showed a very strong correlation between brain and behavior (right panel; r = 0.87). See main text for details of the analysis. A computational notebook for this example is available at https://osf.io/spr9a/.

The problem of multiplicity was recognized very early, and the last 25 years have seen the development of well-established and validated methods for correction of familywise error and false discovery rate in neuroimaging data^45^. However, recent work^44^ has suggested that even some very well-established inferential methods based on spatial extent of activations can produce inflated Type I error rates in certain settings.

There is an ongoing debate between neuroimaging researchers who feel that conventional approaches to multiple comparison correction are too lax and allow too many false positives^46^, and those who feel that thresholds are too conservative, and risk missing most of the interesting effects^47^. In our view, the deeper problem is the inconsistent application of principled correction approaches^48^. Many researchers freely combine different approaches and thresholds in ways that produce a high number of undocumented researcher degrees of freedom^8^, rendering reported p-values uninterpretable.

To assess this more directly, we examined the top 100 results for the Pubmed query (“fMRI” AND brain AND activation NOT review[PT] AND human[MESH] AND english[la]), performed May 23, 2016; of these, 65 reported whole-brain task fMRI results and were available in full text (full list of papers and annotations available at https://osf.io/spr9a/). Only three presented fully uncorrected results, with four others presenting a mixture of corrected and uncorrected results; this suggests that corrections for multiple comparisons are now standard. However, there is evidence that researchers may engage in “method-shopping” for techniques that provide greater sensitivity, at a potential cost of increased error rates. Nine of the 65 papers used the FSL or SPM software packages to perform their primary analysis, but then used the alphasim or 3dClustSim tools from the AFNI software package (7 papers) or other simulation-based approaches (2 papers) to correct for multiple comparisons. This is concerning, because both FSL and SPM offer well-established methods that use Gaussian random field theory or nonparametric analyses to correct for multiple comparisons. Given the substantial degree of extra work (e.g. software installation, file reformatting) involved in using multiple software packages, the use of a different tool raises some concern that this might reflect analytic p-hacking. This concern is further amplified by the finding that until very recently, this AFNI program had slightly inflated inflated Type I error rates^44^. Distressingly, whereas nonparametric (randomization/permutation) methods are known to provide the more accurate control over familywise error rates compared to parametric methods^45,49^ and are applicable for nearly all models, they were not used in any of these papers.

#### Solutions

To balance Type I and Type II error rates in a principled way, we suggest a dual approach of reporting corrected whole-brain results, and (for potential use in later meta-analyses) sharing a copy of the unthresholded statistical map (preferably Z values) through a repository that allows viewing and downloading (such as Neurovault.org^50^). For an example of this practice, see ref. ^51^ and shared data at http://neurovault.org/collections/122/. Any use of non-standard methods for correction of multiple comparisons (for example, using tools from different packages for the main analysis and the multiple comparison correction) should be justified explicitly (and reviewers should demand such justification). Both voxel- and cluster-wise inferences with FWE or FDR correction are suitable to control false positive risk, though cluster-wise and any FDR inferences need to be interpreted with care as they allow more false positive voxels than voxel-wise FWE under typical parameterizations.

Alternatively one can abandon the mass univariate approach altogether. Multivariate methods that treat the entire brain as the measurement (e.g.^52^), and graph-based approaches that integrate information over all edges (e.g.^53^) avoid the multiple testing problem. However, these approaches then make it challenging to understand the involvement of individual voxels/edges in an effect^54^ and raise other interpretation issues.

### Problem: Software errors

As the complexity of a software program increases, the likelihood of undiscovered bugs quickly reaches certainty^55^, which implies that all software used for fMRI analysis is likely to have bugs. Most fMRI researchers use one of several open-source analysis packages for preprocessing and statistical analyses; many additional analyses require custom programs. Because most researchers writing custom code are not trained in software engineering, there is insufficient attention to good software development practices that could help catch and prevent errors. This issue came to the fore recently, when a 15-year-old bug was discovered in the AFNI program 3dClustSim (and the older AlphaSim), which resulted in slightly inflated Type I error rates^44,56^ (the bug was fixed in May 2015). While small in this particular case, the impact of such bugs could be widespread; for example, PubMed Central lists 1362 publications mentioning AlphaSim or 3dClustSim published prior to 2015 (query [("alphasim" OR "3DClustSim") AND 1992:2014[DP]] performed July 14, 2016). Similarly, the analyses presented in a preprint of the present article contained two software errors that led to different results being presented in the final version of the paper. The discovery of these errors led us to perform a code review and to include software tests in order to reduce the likelihood of remaining errors. Even though software errors will happen in commonly used toolboxes as well as in-house code, they are much more likely to be discovered in widely used packages due to scrutiny by many more users. It is very likely that significant bugs exist in custom software built for individual projects, but due to the limited user base those bugs will never be unearthed.

#### Solutions

Researchers should avoid the trap of the “not invented here” philosophy: When the problem at hand can be solved using software tools from a well-established project, these should be chosen instead of re-implementing the same method in custom code. Errors are more likely to be discovered when code has a larger userbase, and larger projects are more likely to follow better software-development practices. Researchers should learn and implement good programming practices, including the judicious use of software testing and validation. Validation methodologies (such as comparing with another existing implementation or using simulated data) should be clearly defined. Custom analysis code should always be shared upon manuscript submission (for an example, see^57^). It may be unrealistic to expect reviewers to evaluate code in addition to the manuscript itself, although this is standard in some journals such as the *Journal of Statistical Software*. However, reviewers should request that the code be made available publicly (so others can evaluate it) and in case of methodological papers that the code is accompanied with a set of automated tests. Finally, researchers need to acquire sufficient training on the implemented analysis methods, in particular so they understand the software’s default parameter values, as well as the assumptions on the data and how to verify those assumptions.

### Problem: Insufficient study reporting

In order to know whether appropriate analyses have been performed, the methods must be reported in sufficient detail. Eight years ago we^58^ published an initial set of guidelines for reporting the methods used in an fMRI study. Unfortunately, reporting standards in the fMRI literature remain poor. Carp^59^ and Guo and colleagues^60^ analyzed 241 and 100 fMRI papers respectively for the reporting of methodological details, and both found that some important analysis details (e.g. interpolation methods, smoothness estimates) were rarely described. Consistent with this, in 22 of the 65 papers discussed above it was impossible to identify exactly which multiple comparison correction technique was used (beyond generic terms such as “cluster-based correction”) because no specific method or citation was provided. The Organization for Human Brain Mapping has recently addressed this issue through its 2015–2016 Committee on Best Practices in Data Analysis and Sharing (COBIDAS), which has issued a new, detailed set of reporting guidelines^61^ (http://www.humanbrainmapping.org/COBIDAS) (see Box 4).

Beyond the description of methods, claims in the neuroimaging literature are often advanced without corresponding statistical support. In particular, failures to observe a significant effect often lead researchers to proclaim the absence of an effect—a dangerous and almost invariably unsupported acceptance of the null hypothesis. “Reverse inference” claims, in which the presence a given pattern of brain activity is taken to imply a specific cognitive process (e.g., “the anterior insula was activated, suggesting that subjects experienced empathy”), are rarely grounded in quantitative evidence^15,62^. Furthermore, claims of “selective” activation in one brain region or experimental condition are often made when activation is statistically significant in one region or condition but not in others—ignoring the fact that “the difference between significant and non-significant is not itself significant”^63^ and in the absence of appropriate tests for statistical interactions^64^.

#### Solutions

Authors should follow accepted standards for reporting methods (such as the COBIDAS standard for MRI studies), and journals should require adherence to these standards. Every major claim in a paper should be directly supported by appropriate statistical evidence, including specific tests for significance across conditions and relevant tests for interactions. Because the computer code is often necessary to understand exactly how a dataset has been analyzed, releasing the analysis code is particularly useful and should be adopted as a standard practice.

### Problem: Lack of independent replications

There are surprisingly few examples of direct replication in the field of neuroimaging, likely reflecting both the expense of fMRI studies along with the emphasis of most top journals on novelty rather than informativeness. While there are many basic results that are clearly replicable (e.g. presence of face-selective vs. scene-selective activation in ventral temporal cortex, or systematic correlations within functional networks during the resting state), the replicability of more subtle and less neurobiologically established effects (such as group differences and between-subject correlations) is nowhere near as certain. One study^65,66^ attempted to replicate 17 studies that had previously found associations between brain structure and behaviour. Only one of the 17 replication attempts showed stronger evidence for an effect as large the original effect size rather than for a null effect, and 8 out of 17 showed stronger evidence for a null effect. This suggests that replicability of neuroimaging findings (particularly brain-behavior correlations) may be exceedingly low, similar to recent findings in other areas of science such as cancer biology^67^ and psychology^68^.

It is worth noting that even though the cost of conducting an new fMRI experiment is a factor limiting the feasibility of replications studies there are many findings that can be replicated using publicly available data. Resources such as FCP-INDI^26^, CoRR^69^, OpenfMRI^70^, or the Human Connectome Project^19^ provide MRI data suitable to attempt to replicate many previously reported findings. They can also be used to answer questions about sensitivity of of a particular finding to the data analysis tools used^35^. However, even in the cases when a replications are possible using publicly available data they are still rare and far apart, because the academic community puts much bigger emphasis on novelty of findings rather than their replicability.

#### Solutions

The neuroimaging community should acknowledge replication reports as scientifically important research outcomes that are essential in advancing knowledge. One such attempt is the OHBM Replication Award to be awarded in 2017 for the best neuroimaging replication study in the previous year. In addition, in case of especially surprising findings (for example extra-sensory perception), findings that could have influence on public health policy or medical treatment decisions, or findings that could be tested using data from another existing dataset, reviewers should consider requesting replication of the finding by the group before accepting the manuscript.

### Towards the neuroimaging paper of the future

In the foregoing we have outlined a number of problems with current practice and made suggestions for improvements. Here we outline what we would like to see in the “neuroimaging paper of the future”, inspired by related work in the geosciences^71^.

#### Planning

The sample size for the study would be determined in advance using formal statistical power analysis. The entire analysis plan would be formally pre-registered, including inclusion/exclusion criteria, software workflows including contrasts and multiple comparison methods, and specific definitions for all planned regions of interest.

#### Implementation

All code for data collection and analysis would be stored in a version control system, and would include software tests to detect common problems. The repository would use a continuous integration system to ensure that each revision of the code passes the appropriate software tests. The entire analysis workflow (including both successful and failed analyses) would be completely automated within a workflow engine and packaged within a software container or virtual machine in order to ensure computational reproducibility. All datasets and results would be assigned version numbers in order to allow explicit tracking of provenance. Automated quality control would assess the analysis at each stage to detect potential errors.

#### Validation

For empirical papers, all exploratory results would be validated against an independent validation dataset that was not examined prior to validation. For methodological papers, the approach would follow best practices for reducing overly optimistic results^72^. Any new method would be validated against benchmark datasets and compared to other state-of-the-art methods.

#### Dissemination

All results would be clearly marked as either hypothesis driven (with a link to the appropriate pre-registration) or exploratory. All analyses performed on the dataset (including those that failed) would be reported. The paper would be written using a literate programming technique, in which the code for figure generation is embedded within the paper and the image data depicted in figures is transparently accessible. The paper would be distributed along with the full codebase to perform the analyses and the data necessary to reproduce the analyses, preferably in a container or virtual machine to allow direct reproducibility. Both unthresholded statistical maps and the raw data would be shared via appropriate community repositories, and the shared raw data would be formatted according to a community standard, such as the Brain Imaging Data Structure (BIDS)^73^, and annotated using an appropriate ontology to allow automated meta-analysis.

### Conclusion

We have outlined what we see as a set of problems with neuroimaging methodology and reporting and approaches to address them. It is likely that the reproducibility of neuroimaging research is no better than many other fields, where it has been shown to be surprisingly low. Given the substantial amount of research funds currently invested in neuroimaging research, we believe that it is essential that the field address the issues raised here, so as to ensure that public funds are spent effectively and in ways that advance our understanding of the human brain. We have also laid out what we see as a roadmap for how neuroimaging researchers can overcome these problems, laying the groundwork for a scientific future that is transparent and reproducible.

## Further information

Fmripower: fmripower.org

Human Connectome Project: https://www.humanconnectome.org/

Organisation for Human Brain Mapping (OHBM): www.humanbrainmapping.org

NeuroPower: neuropowertools.org Neurosynth: http://neurosynth.org/

Neurovault: http://neurovault.org/

#### Box 1: Lessons from Genetics

The study of genetic influences on complex traits has been transformed by the advent of whole genome methods, and the subsequent use of stringent statistical criteria, independent replication, large collaborative consortia, and complete reporting of statistical results. Previously, “candidate” genes would be selected on the basis of known or presumed biology, and a handful of variants genotyped (many of which would go unreported) and tested in small studies. An enormous literature proliferated, but these findings generally failed to replicate^74^. The transformation brought about by genome wide association studies applied in very large populations was necessitated by the stringent statistical significance criteria required by simultaneous testing of several hundred thousand genetic loci, and an emerging awareness that any effects of common genetic variants generally are very small (#x003C;1% phenotypic variance). To realise the very large sample sizes required, large-scale collaboration and data sharing was embraced by the genetics community. The resulting cultural shift has rapidly transformed our understanding of the genetic architecture of complex traits, and in a few years produced many hundred more reproducible findings than in the previous fifteen years^75^! Routine sharing of single nucleotide polymorphism (SNP)-level statistical results has facilitated routine use of meta-analysis, as well as the development of novel methods of secondary analysis^76^.

This relatively rosy picture contrasts markedly with the situation in “imaging genetics”--a burgeoning field that has yet to embrace the standards commonly followed in the broader genetics literature, and which remains largely focused on individual candidate gene association studies, which are characterized by numerous researcher degrees of freedom. To illustrate, we examined the first 50 abstracts matching a PubMed search for “fMRI” and “genetics” (excluding reviews, studies of genetic disorders, and nonhuman studies) which included a genetic association analysis (for list of search results, see https://osf.io/spr9a/). Of these, the vast majority (43/50) reported analysis of a single or small number (5 or fewer) of candidate genes; only 2/50 reported a genome-wide analysis, with the rest reporting analyses using biologically inspired gene sets (3/50) or polygenic risk scores (2/50). Recent empirical evidence also casts doubt on the validity of candidate gene associations in imaging genomics. A large genome-wide association study of whole-brain and hippocampal volume^77^ identified two genetic associations that were both replicated across two large samples each containing more than 10,000 individuals. Strikingly, analysis of a set of candidate genes previously reported in the literature showed no evidence for any association in this very well-powered study^77^. The lessons for imaging from genome-wide association studies more generally seems clear: associations of common genetic variants with complex behavioual phenotypes are generally very small (<1% of phenotypic variance) and thus require large, homogeneous samples to be able to identify them robustly. As the prior odds for association of any given genetic variant with novel imaging phenotypes generally are low, and given the large number of variants simultaneously tested in a genome-wide association study (necessitating a corrected P-value threshold of ~10^−8^), adequate statistical power can only be achieved by using sample sizes in the many thousands to tens of thousands. Finally, results need to be replicated to ensure robust discoveries.

#### Box 2: Effect-size estimates for common neuroimaging experimental paradigms

The aim of this analysis is to estimate the magnitude of typical effect sizes of blood oxygen level-dependent changes in fMRI signal associated with common psychological paradigms. We focus on four experiments administered by the Human Connectome Project (HCP)^78^: an emotion task, gambling task, working memory task and motor task (detailed below). We chose data from the HCP for its large sample size, which allows computation of stable effect size estimates, and its diverse set of activation tasks. The data and code used for this analysis are available at https://osf.io/spr9a/.

Briefly, the processing of data from the Human Connectome Project was carried out in 4 main steps:

1. **Subject Selection:** The analyses are performed on the 500 subjects release of the HCP data, freely available at www.humanconnectome.org. We selected 186 independent subjects from the HCP data on the bases that (1) all subjects have results for all four of the tasks and (2) there are no genetically related subjects in the analysis.
2. **Group Analyses:** The first-level analyses, which summarise the relation between the experimental design and the measured time series for each subject, were obtained from the Human Connectome Project. The processing and analysis pipelines for these analyses are shared together with the data. Here we perform second-level analyses — that is, an assessment of the average effect of the task on BOLD signal over subjects — using the FSL program flame1^17^ which performs a linear mixed-effects regression at each voxel, using generalized least squares with a local estimate of random effects variance. This analysis averages over subjects, while separating within-subject and between-subject variability to ensure control of unobserved heterogeneity. The specific contrasts that were tested are:

- Motor: average BOLD response for tongue, hand and foot movements versus rest
- Emotion: viewing faces with a fearful expression versus viewing neutral faces
- Gambling: monetary reward versus punishment
- Working memory: a contrast between conditions in which the participants indicate whether the current stimulus matches the one from 2 trials earlier (“2-back”), versus a condition where the participants indicate whether the current stimulus matches a specific target (“0-back”)
3. **Create Masks:** The masks used for the analyses are the intersections of anatomical and *a priori* functional masks for each contrast. The rationale behind this is to find effect sizes in regions that are functionally related to the task, but restricted to certain anatomical regions.
  - Functional: We created masks using www.neurosynth.org^79^. To do this, we performed forward inference meta-analysis using the respective search terms "Motor","Emotion","Gambling","Working memory" for each of the tasks, with false discovery rate (FDR) control at 0.01, the default threshold on neurosynth. The resulting mask identifies voxels consistently found to be activated in studies that mention each of the search terms in their abstract.
  - Anatomical: We have used Harvard-Oxford probabilistic atlas^22^ at p>0. Regions were chosen for each task based on the published a priori hypothesized regions from the HCP^23^. The size of the masks was assessed by the number of voxels in the mask.
4. **Compute Effect Size:** The intersection masks created above were used to isolate the regions of interest in the second-level-analysed BOLD signal data. From these mask-isolated data sets, the size of the task-related effect (Cohen’s D) were computed for each relevant region(see Figure B2 below). FSL’s Featquery computes for each voxel the % BOLD change in the data within the masks.

| Task | Anatomical Mask |
| --- | --- |
| Motor | • Precentral gyrus<br>• Supplementary motor cortex<br>• Left putamen<br>• Right putamen |
| Working memory | Middle frontal gyrus |
| Emotion | • Left amygdala<br>• Right amygdala |
| Gambling | • Left accumbens<br>• Right accumbens |

**Figure B2:**
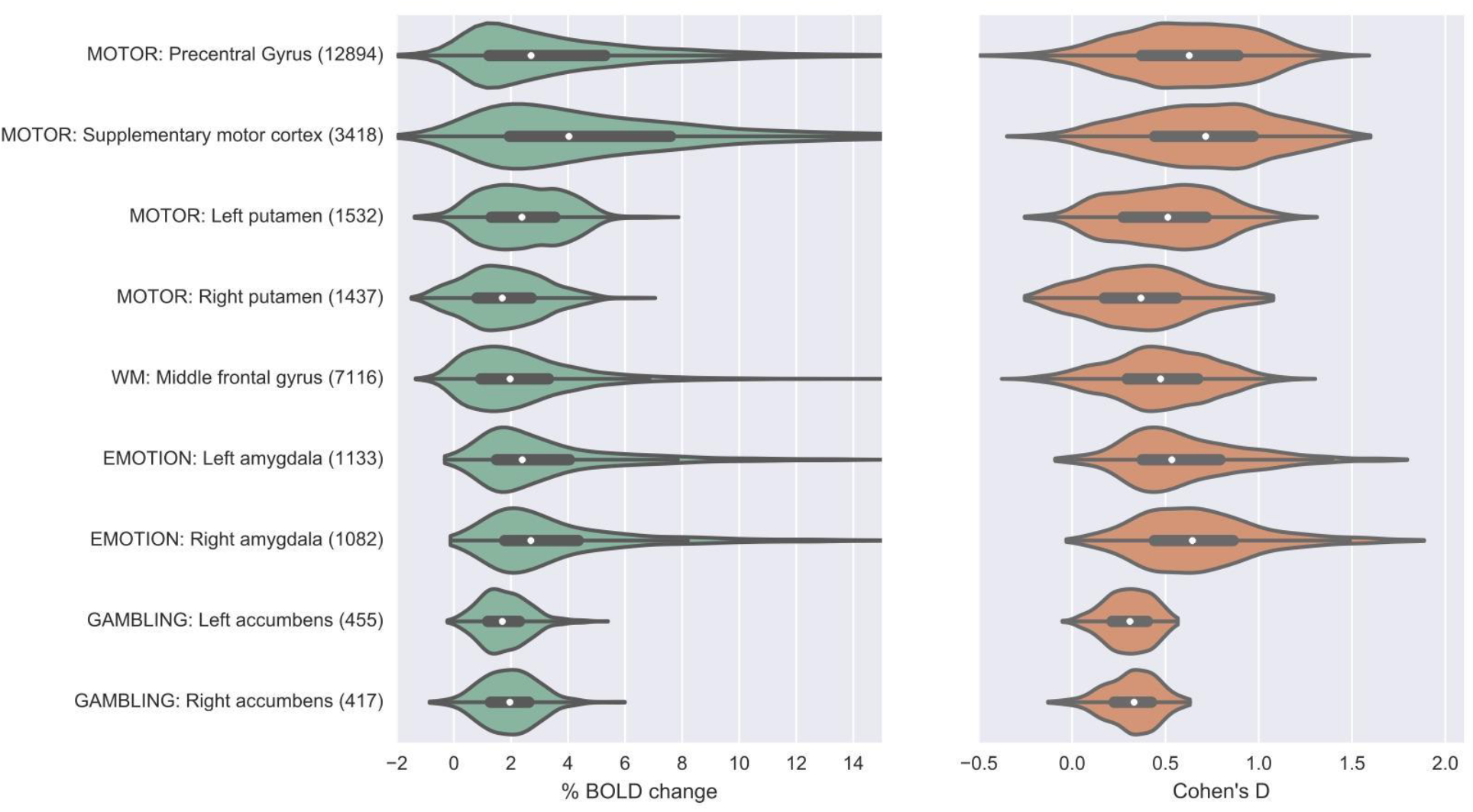
The distributions of the observed effect size estimates and BOLD signal change estimates for common experimental paradigms, across voxels within each ROI (number in parentheses denotes the number of voxels in the ROI). The boxplot inside the violins represent the interquartile range (first quartile to third quartile) and the white dot shows the median value.

#### Box 3: Flexibility in fMRI data analysis

In the early days of fMRI analysis, it was rare to find two laboratories that used the same software to analyze their data, with most using locally-developed custom software. Over time, a small number of open-source analysis packages have gained prominence (SPM, FSL, and AFNI being the most common), and now most laboratories use one of these packages for their primary data processing and analysis. Within each of these packages, there is a great deal of flexibility in how data are analyzed; in some cases there are clear best practices, but in other cases there is no consensus regarding the optimal approach. This leads to a multiplicity of analysis options. In Table B1 we outline some of the major choices involved in performing analyses using one of the common software packages (FSL). Even for this non-exhaustive list from a single analysis package, the number of possible analysis workflows exceeds the number of papers that have been published on fMRI since its inception more than two decades ago!

It is possible that many of these alternative pipelines could lead to very similar results, though the analyses of Carp^35^ suggest that many of them may lead to significant heterogeneity in the results. In addition, there is evidence that choices of preprocessing parameters may interact with the statistical modeling approach (e.g., interactions between head motion modeling and physiological noise correction), and that the optimal preprocessing pipeline may differ across subjects (e.g. interacting with the amount of head motion)^42^.

**Table B3:** A non-exhaustive list of data processing/analysis options available within the FSL software package, enumerating a total of 69,120 different possible workflows.

| Processing step | Reason | Options [suboptions] | Number of plausible options |
| --- | --- | --- | --- |
| Motion correction | Correct for head motion during scanning | Interpolation [linear vs. sinc]<br>Reference volume [single vs. mean] | 4 |
| Slice timing correction | Correct for differences in acquisition timing of different slices | No/before motion correction/after motion correction | 3 |
| Field map correction | Correct for distortion due to magnetic susceptibility | Yes/No | 2 |
| Spatial smoothing | Increase SNR for larger activations and ensure assumptions of Gaussian random field theory | FWHM [4/6/8 mm] | 3 |
| Spatial normalization | Warp individual brain to match a group template | Method [linear/nonlinear] | 2 |
| High pass filter | Remove low-frequency nuisance signals from data | Frequency cutoff [100 secs, 120 secs] | 2 |
| Head motion regressors | Remove remaining signals due to head motion via statistical model | Yes/No<br>If Yes: 6/12/24 parameters or single timepoint “scrubbing” regressors | 5 |
| Hemodynamic response | Account for delayed nature of hemodynamic response to neuronal activity | Basis function [single-gamma, double-gamma]<br>Derivatives [none/shift/dispersion] | 6 |
| Temporal autocorrelation model | Model for the temporal autocorrelation inherent in fMRI signals. | Yes/no | 2 |
| Multiple comparison correction | Correct for large number of comparisons across the brain | Voxel-based GRF, Cluster-based GRF, FDR, nonparametric | 4 |
| **Total possible workflows** |  |  | **69,120** |

#### Box 4: Guidelines for transparent methods reporting in neuroimaging

The OHBM COBIDAS report provides a set of best practices for reporting and conducting studies using MRI. It divides practice into seven categories, and provides detailed checklists that can be consulted when planning, analyzing and writing up a study. The table below lists these categories with summaries of the topics covered in the checklists.

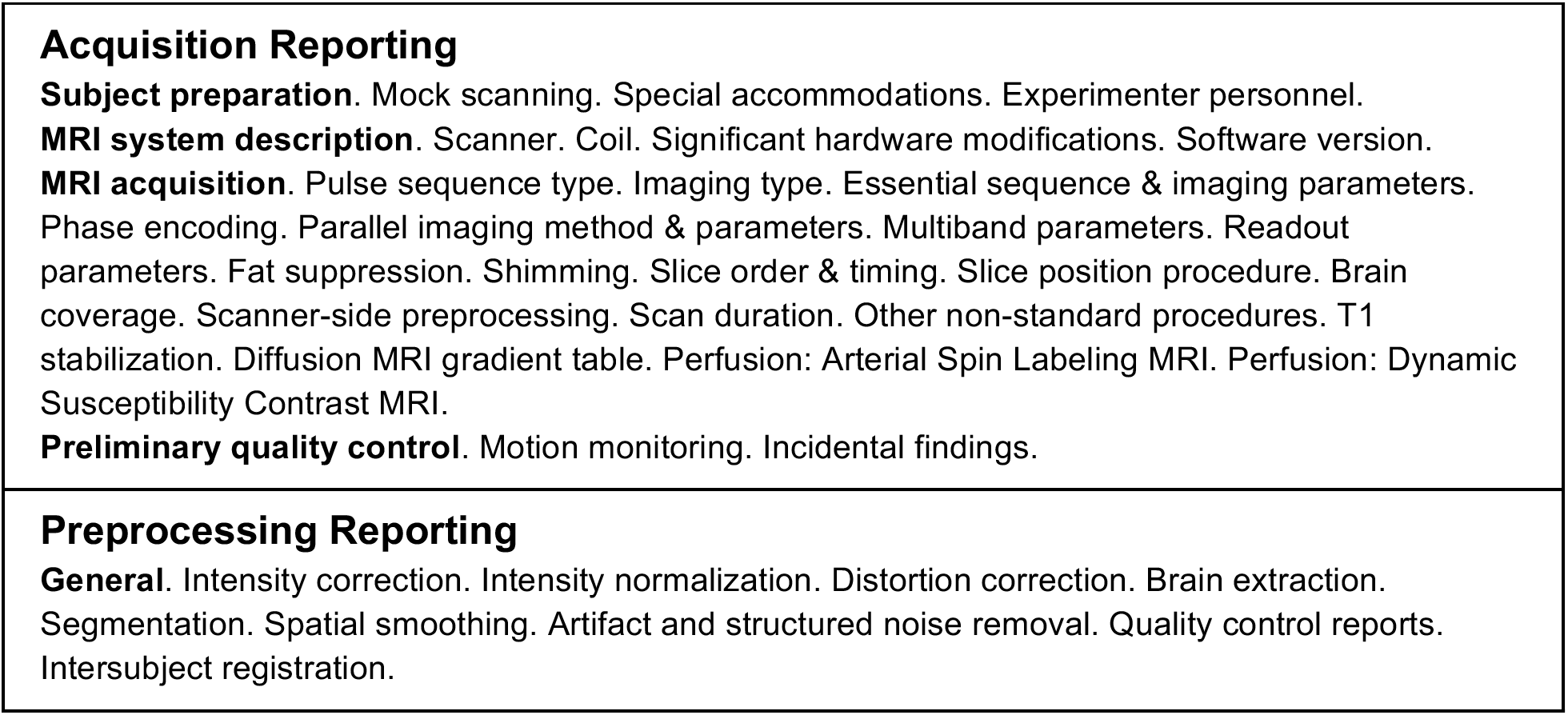
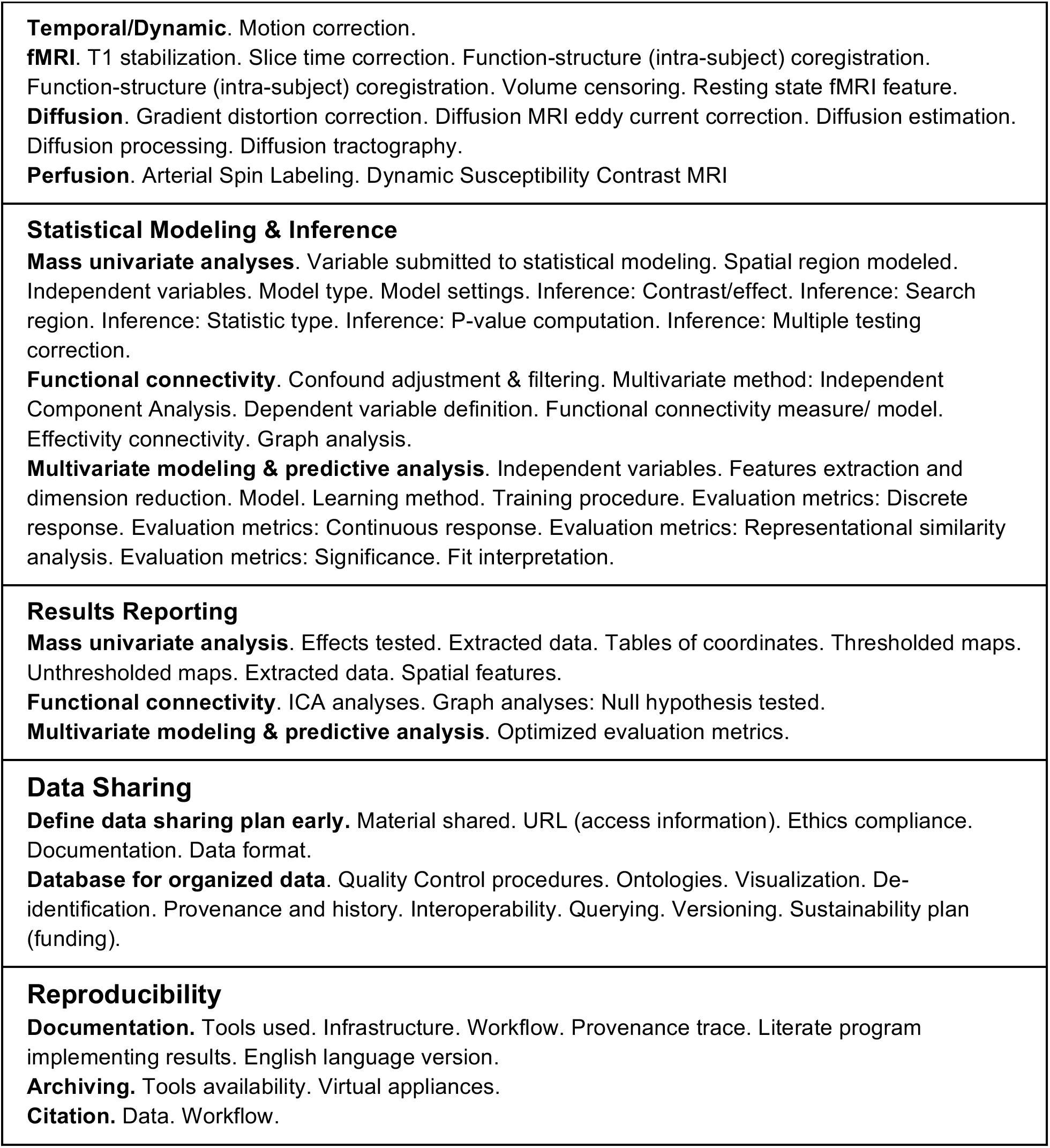

## Acknowledgements

RP, JD, JBP and KG are supported by the Laura and John Arnold Foundation. MRM is supported by the MRC (MC UU 12013/6) and a member of the UK Centre for Tobacco and Alcohol Studies, a UK Clinical Research Council Public Health Research: Centre of Excellence. Funding from British Heart Foundation, Cancer Research UK, Economic and Social Research Council, Medical Research Council, and the National Institute for Health Research, under the auspices of the UK Clinical Research Collaboration, is gratefully acknowledged. CIB is supported by the Intramural Research Program of NIMH (ZIA-MH 002909). TY is supported by NIMH (R01MH096906). PMM gratefully acknowledges personal support from the Edmond J. Safra Foundation and Lily Safra and research support from the Medical Research Council, the Imperial College Healthcare Trust Biomedical Research Centre and the Imperial EPSRC Mathematics in Healthcare Centre. TEN is supported by the Wellcome Trust (100309/Z/12/Z). JBP is supported by NIH-NIBIB P41-EB019936 and by NIH-NIDA U24-038653. Data were provided [in part] by the Human Connectome Project, WU-Minn Consortium (Principal Investigators: David Van Essen and Kamil Ugurbil; 1U54MH091657) funded by the 16 Institutes and Centers of the US National Institutes of Health (NIH) that support the NIH Blueprint for Neuroscience Research; and by the McDonnell Center for Systems Neuroscience at Washington University. Thanks to Joe Wexler for performing annotation of Neurosynth data, Sean David for providing sample size data, and Robert Cox and Paul Taylor for helpful comments on a draft of the manuscript.

## Supplementary Materials

**Figure S1:**
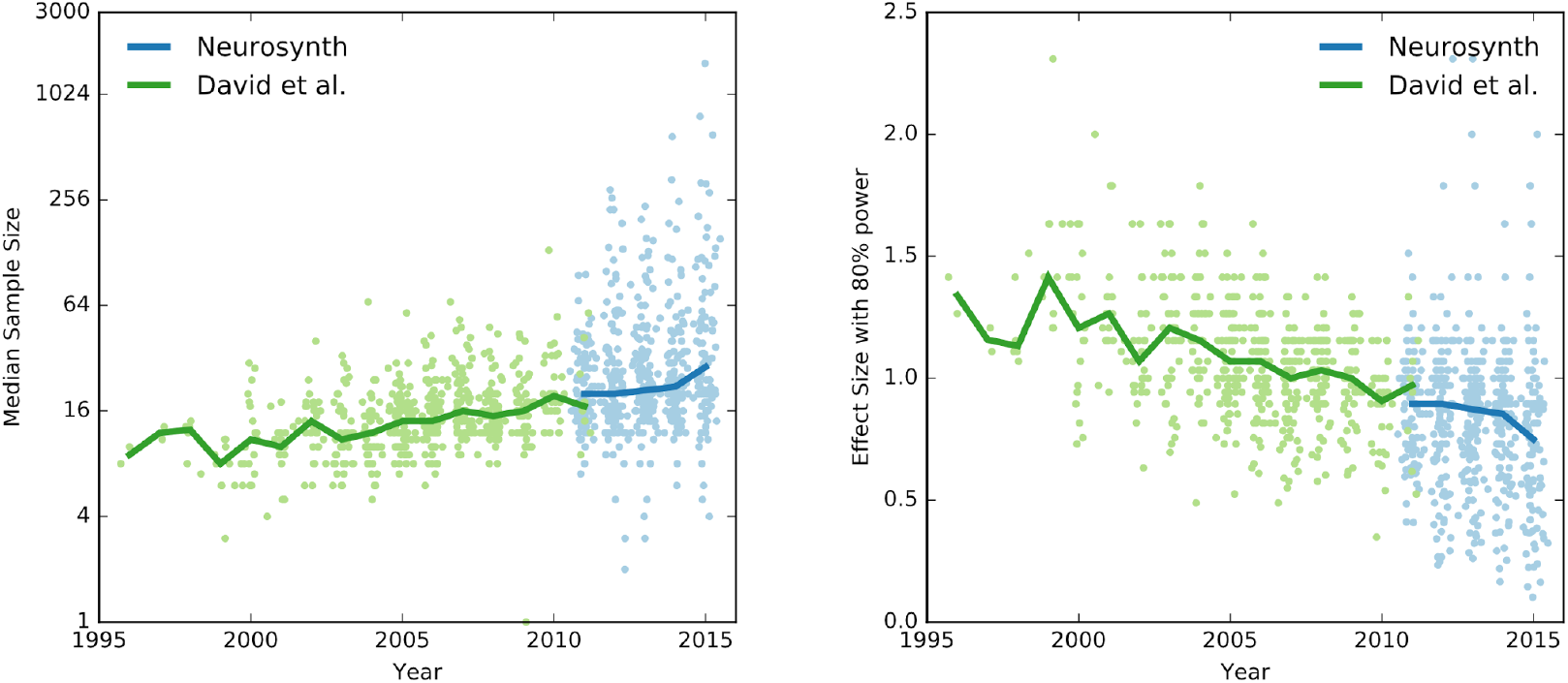
A depiction of the data from Figure 1 showing all data points. Sample sizes are shown on a log scale.

